# Structure of mechanically activated ion channel OSCA2.3 reveals mobile elements in the transmembrane domain

**DOI:** 10.1101/2023.06.15.545135

**Authors:** Sebastian Jojoa-Cruz, Batuujin Burendei, Wen-Hsin Lee, Andrew B. Ward

**Affiliations:** Department of Integrative Structural and Computational Biology, Scripps Research, La Jolla, California 92037, USA

**Keywords:** OSCA/TMEM63, mechanosensation, mechanically activated ion channels, cryo-EM, peptidisc

## Abstract

Members of the OSCA/TMEM63 are mechanically activated ion channels and structures of some OSCA members have revealed the architecture of these channels and structural features that are potentially involved in mechanosensation. However, these structures are all in a similar state and information about the motion of different elements of the structure is limited, preventing a deeper understanding of how these channels work. Here, we used cryo-electron microscopy to determine high resolution structures of *Arabidopsis thaliana* OSCA1.2 and OSCA2.3 in peptidiscs. The structure of OSCA1.2 resembles previous structures of the same protein in different environments. Yet, in OSCA2.3 the TM6a-TM7 linker constricts the pore on its cytoplasmic side, revealing conformational heterogeneity within the OSCA family. Furthermore, coevolutionary sequence analysis uncovered a conserved interaction between TM6a-TM7 linker and the Beam-Like Domain. Our results support the involvement of TM6a-TM7 in mechanosensation and potentially in the diverse response of OSCA channels to mechanical stimuli.

## Introduction

Mechanically activated (MA) ion channels allow organisms to sense external and internal mechanical cues^1,2^, and correct interpretation of these cues is essential for survival. In plants, MA ion channels have been linked to roles like pollen fertility^3^, induced cell death^4^, cell swelling in seedlings^5^, and plant stomatal immunity^6^. OSCA/TMEM63 is a family of MA ion channels found to be conserved across different eukaryotic clades^7^ and plant OSCAs have been found to play a crucial role in osmotic stress response^8-13^. Climate change and the accompanying increase in severity of abiotic stress such as drought and salinity, which cause osmotic stress, have brought dramatic negative effects on crop yields^14^. Transgenic expression of OSCAs has led to improved osmotic stress tolerance^10,11,15^. Thus, understanding how these mechanosensors operate may provide insights into coping with changing and increasingly harsh environmental conditions.

The OSCA family in plants is diverse, usually with several OSCA genes per genome divided in clades 1 through 4^8,13,15-17^. For instance, *Arabidopsis thaliana* (At) has 15 genes belonging to this family^8,16^, and such variety hints at broad functional diversity^1,18^. Accordingly, *in vitro* characterization of these channels has shown differential response to mechanical stimuli^7^, while *in vivo* experiments indicate involvement of select OSCAs in biological processes^6,8,10^. But it remains to be seen if such differences correlate with structural differences at the protein level. Thus far, high-resolution structures of OSCAs are limited to two members of clade 1, OSCA1.1^19^ and OSCA1.2^20-22^ from At with 85% sequence identity.

OSCAs are structurally homologous to TMEM16s and consist of a dimeric channel with a pore on each protomer that is accessible to membrane lipids through a side opening on its intracellular half^19-22^. The intracellular domain is the sole responsible of channel oligomerization and presents two membrane-interacting elements proposed to be involved in mechanosensation: the amphipathic helix and the Beam-Like Domain (BLD). Currently available structures were solved in a closed state^19-22^, hindering a mechanistic understanding of OSCA mechanosensation.

To further explore the differences within OSCA clades we determined the structure of OSCA2.3, which exhibits single-channel currents in the sub-picoampere range and distinct electrophysiological properties compared to other characterized OSCAs including slower activation and inactivation kinetics, ^7^. This high-resolution cryo-electron microscopy (cryo-EM) structure of AtOSCA2.3 was solved in peptidiscs^23^. The conformation of OSCA2.3 relative to previous OSCA structures reveals the TM6a-TM7 linker as a mobile element that may play a role in transition between states of the channel, including its involvement as a potential inactivation gate. We also solved the cryo-EM structure of AtOSCA1.2 in peptidiscs as a comparator and demonstrate that protein interactions with the peptidisc did not induce any meaningful structural differences.

## Results

### Structure of OSCA2.3 and OSCA1.2 in peptidiscs

Full-length cDNA of AtOSCA2.3 (UniProt ID:Q8GUH7) was genetically fused with C-terminal EGFP and expressed in mammalian cells. Protein was solubilized with DDM+CHS, purified, reconstituted into peptidiscs^23^, and imaged by cryo-EM. The resulting 2.7Å map of OSCA2.3 enabled modelling of the majority of the protein (504 out of 703 residues) (Fig. 1a-c, Table 1, Supplementary Fig. 1-3). The general architecture of OSCA2.3 matched those of previous OSCA structures solved in detergent or nanodiscs^19-22^, with two protomers of 11 transmembrane helices (TMs) each, an intracellular domain that makes the dimerization domain and presence of the putative mechanosensitive amphipathic helix. However, the transmembrane domain (TMD) exhibited significant movements in the position of the pore-lining TMs relative to previous OSCAs^19-22^. Despite several attempts, structures of OSCA2.3 in detergent/nanodiscs could not be obtained. Thus, to rule out the peptidisc as the cause of these structural differences observed in OSCA2.3, we solved a 2.8Å peptidisc-reconstituted structure of AtOSCA1.2, which has previously been solved in different environments^20,21^ (Fig. 1d-f, Table 1, Supplementary Fig. 1,2). The overall architecture of OSCA1.2 protomer and the pore profile was nearly identical between the different membrane mimetic environments (Supplementary Fig. 4). There was however a modest global shift (∼3.5 Å) in the position of protomers relative to each other (Supplementary Fig. 4). For all future comparisons we refer to the structure of OSCA1.2 in peptidiscs unless otherwise specified. These results suggest that the changes in the TMD of OSCA2.3 are not induced by interactions with the peptidisc. One notable difference between the two structures in peptidiscs is the very well-defined density for four peptidisc peptides per protomer in OSCA1.2 map (Fig. 1d, e). To our knowledge, this constitutes the first time that side chain assignment has been possible for these peptides in a cryo-EM map.

**Table 1.**
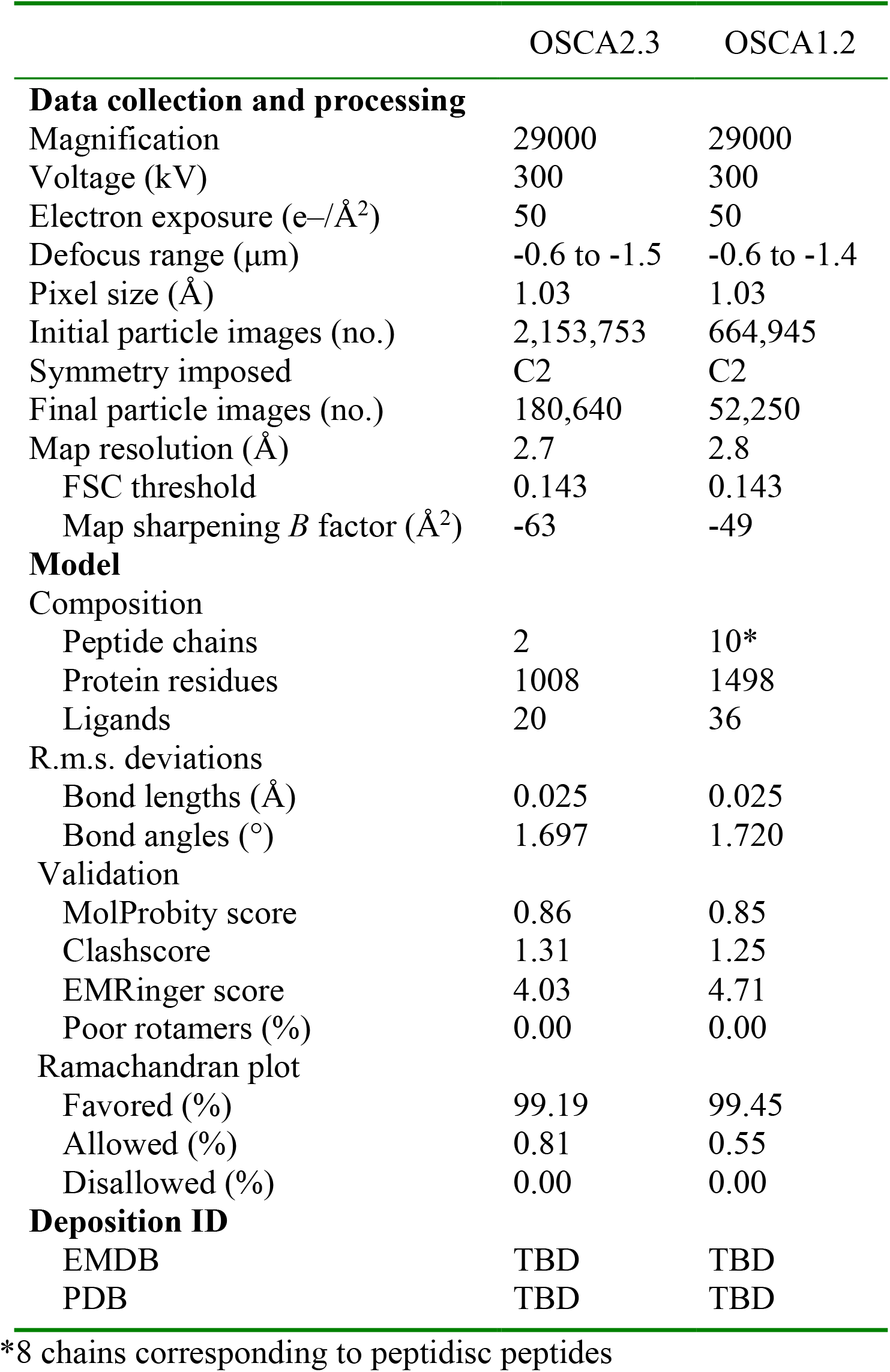
Data collection, processing, model refinement and validation.

|  | OSCA2.3 | OSCA1.2 |
| --- | --- | --- |
| **Data collection and processing** |  |  |
| Magnification | 29000 | 29000 |
| Voltage (kV) | 300 | 300 |
| Electron exposure (e-/Å<sup>2</sup>) | 50 | 50 |
| Defocus range (µm) | -0.6 to -1.5 | -0.6 to -1.4 |
| Pixel size (Å) | 1.03 | 1.03 |
| Initial particle images (no.) | 2,153,753 | 664,945 |
| Symmetry imposed | C2 | C2 |
| Final particle images (no.) | 180,640 | 52,250 |
| Map resolution (Å) | 2.7 | 2.8 |
| FSC threshold | 0.143 | 0.143 |
| Map sharpening *B* factor (Å<sup>2</sup>) | -63 | -49 |
| **Model** |  |  |
| Composition |  |  |
| Peptide chains | 2 | 10* |
| Protein residues | 1008 | 1498 |
| Ligands | 20 | 36 |
| R.m.s. deviations |  |  |
| Bond lengths (Å) | 0.025 | 0.025 |
| Bond angles (°) | 1.697 | 1.720 |
| Validation |  |  |
| MolProbity score | 0.86 | 0.85 |
| Clashscore | 1.31 | 1.25 |
| EMRinger score | 4.03 | 4.71 |
| Poor rotamers (%) | 0.00 | 0.00 |
| Ramachandran plot |  |  |
| Favored (%) | 99.19 | 99.45 |
| Allowed (%) | 0.81 | 0.55 |
| Disallowed (%) | 0.00 | 0.00 |
| **Deposition ID** |  |  |
| EMDB | TBD | TBD |
| PDB | TBD | TBD |

\*8 chains corresponding to peptidisc peptides

**Figure 1.**
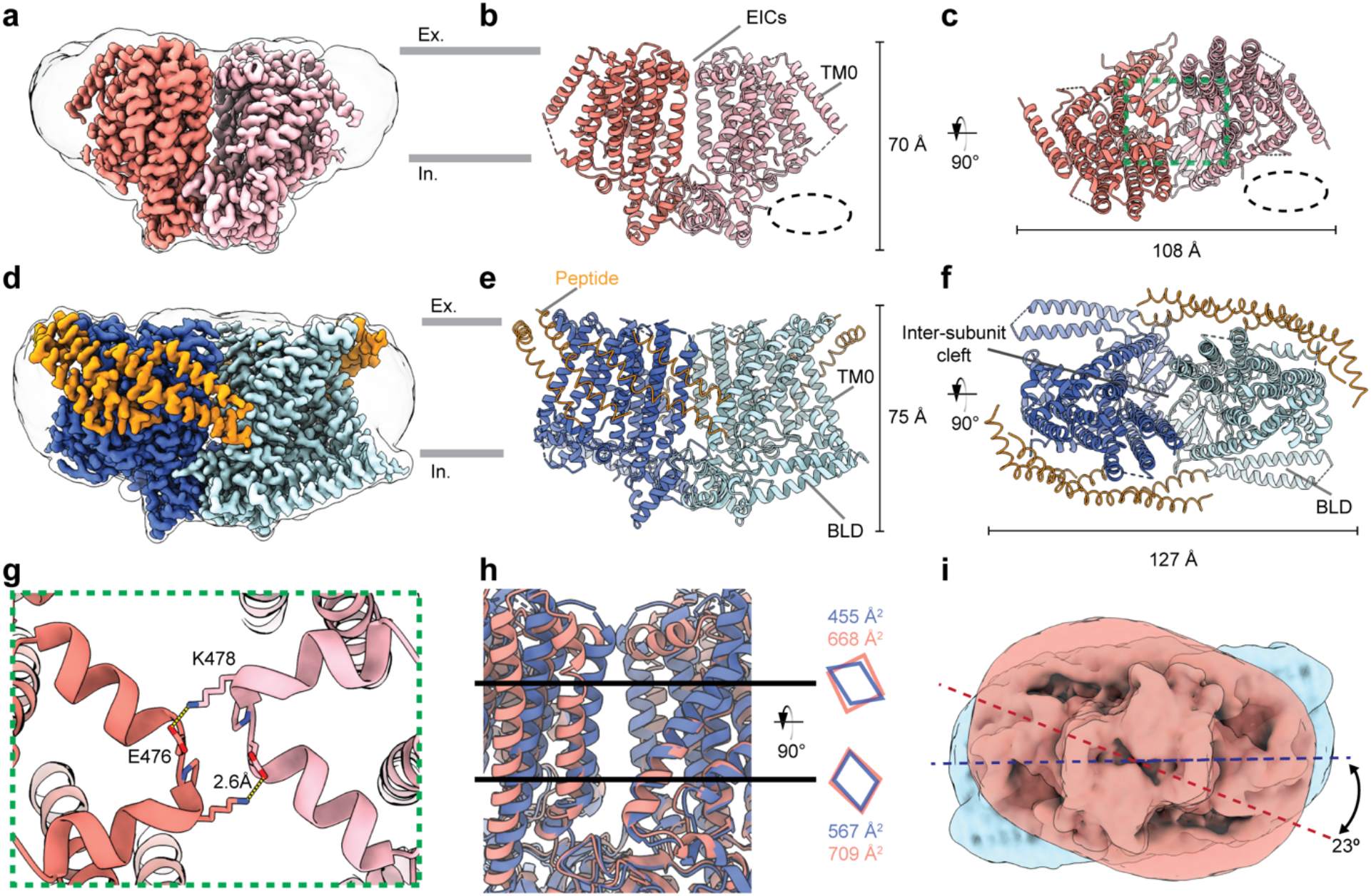
Structure of OSCA2.3 and OSCA1.2 in peptidiscs. DeepEMhancer maps (**a, d**), side (**b, e**) and top (**c, f**) views of models of OSCA2.3 and OSCA1.2, respectively. Ex: Extracellular, In: Intracellular. **g**, Close-up view from **c** displaying the EICs. **h**, Right: Close-up view of the inter-subunit cleft of OSCA2.3 (salmon) and OSCA1.2 (blue). Black lines represent the approximate height at which cross-sectional area was measured. Left: Representation of the shape of the cross sections and measurements of the area occupied. **i**, Superposition of gaussian-filtered unsharpened maps showing difference in overall peptidisc shape of OSCA2.3 (red) and OSCA1.2 (blue).

A peculiar characteristic of the previous OSCA structures is the lack of inter-subunit contacts in the TMD and extracellular side, which causes the formation of an inter-subunit cleft that is filled by lipids^19-22^ (Fig. 1f). In the case of OSCA2.3, two extracellular inter-subunit contacts (EICs) are formed by the TM5-TM6 loop (Fig. 1g). Each protomer contributes residues E476 and K478 to form a salt bridge with the opposite residue of the complementary subunit, These EICs are not present in previous OSCA structures^19-22^.

Despite these interactions, the inter-subunit cleft remains present in the TMD. In fact, the cross-sectional area occupied by the cleft near its bottom and top is larger in OSCA2.3 than OSCA1.2 (Fig. 1h). Several lipid-like densities are visible in the cleft of both OSCA structures (Supplementary Fig. 2b, d). For OSCA2.3, a couple of these densities have been assigned as cholesterol, which could impact the stiffness of the membrane and influence the mechanosensitive activity of the channel, as seen for MA channel Piezo1^24^.

The BLD is composed of two intracellular helices that run parallel to the membrane and the membrane hook, a hydrophobic linker which re-inserts itself into the membrane^20-22^. Based on the uniqueness of this feature and its close interaction with the membrane lipids, it has been proposed to be involved in mechanosensation^20-22^. Unlike structures of OSCA1.1^19^, OSCA1.2^20-22^, and OSCA3.1^19^, the BLD of OSCA2.3 is completely unresolved in the cryo-EM map, consistent with a high degree of flexibility (Fig. 1a-f). Previously, the GXXGXXG motif (with X usually representing a hydrophobic residue) in the hook was identified in AtOSCAs of clade 1 and 3 and OSCA2.3^21^ (Supplementary Fig. 3). However, the BLD was also unresolved in low-resolution map of OSCA2.2, which lacks this motif^21^. These results suggest that other factors besides the presence of this motif may determine the stability of the BLD. Notably, the presence or absence of a well-ordered BLD seems to affect the disposition of the particle on the intracellular side of the membrane (Fig. 1c,f,i).

### TM6a-TM7 linker narrows the pore pathway of OSCA2.3 towards its intracellular end

The pore profile of OSCA2.3 differs from OSCA1.2 and previous OSCA structures, at both its extracellular and intracellular openings. Towards the extracellular side, the most pronounced difference is a kink in the OSCA2.3 TM3 α-helix, which causes the end of TM3 to move away from the TMD (Fig. 2a). This conformational change is similar to that recently observed in cryo-EM structures of lipid-scramblase TMEM16F^25^, a member of the TMEM16 family, which are structurally homologous to OSCAs. In the presence of calcium, the activating point mutation F518H in TMEM16F results in the movement of the extracellular ends of helices TM3 and TM4 (Fig. 2b), such that the pore opens substantially^25^ (Fig. 2e,f). Likewise, the extracellular region of OSCA2.3 shows an opening of the pore caused by the kink, located at a similar position on TM3 as the kink in TMEM16F (Fig. 2b,d,f). Immediately below the extracellular opening, the pore of OSCA2.3 contains a hydrophobic constriction of similar width to the corresponding hydrophobic gate of OSCA1.2 in peptidisc (Fig. 2f). At the cytoplasmic end, the position of TM6a-TM7 linker closes access of the pore of OSCA2.3 to the intracellular vestibule present in OSCA1.2, making the pore largely hydrophobic throughout the extent of the TMD (Fig. 2c,d,f). This linker comprises two helical features, TM6b and Intracellular Loop 4 Helix (IL4H), and the loop that connects to TM7. These features are located towards the cytoplasmic end of the pore in OSCA1.2 but are closer to the TMD in OSCA2.3. The TM6a-TM7 linker of OSCA2.3 decreases the width of the pore to a minimum Van der Waals radius of ∼0.4 Å at approximately 25 Å below the hydrophobic gate, creating a secondary constriction in the permeation pathway (Fig. 2d,f). The conserved residue E510 (E531 in OSCA1.2) of TM6a is expected to contribute to the permeation pathway of OSCAs based on its location and effect on single channel conductance in AtOSCA1.2^20^. However, TM6a-TM7 linker of OSCA2.3 prevents access of E510 to the pore (Fig. 2d,f). In OSCA1.2, TM5 and TM6a π-helical turns are located near the pore constriction and have been suggested to play a role in channel gating^20,22^. In OSCA2.3, only the π-helical turn in TM5 remains, while in TM6a, residue W504 (equivalent of W525 in OSCA1.2) has straightened into a continuous α-helix (Fig. 2c,d). The retention of the extracellular hydrophobic gate and further blockage of the pore by TM6a-TM7 linker indicates that the structure most likely corresponds to a non-conductive, closed state.

**Figure 2.**
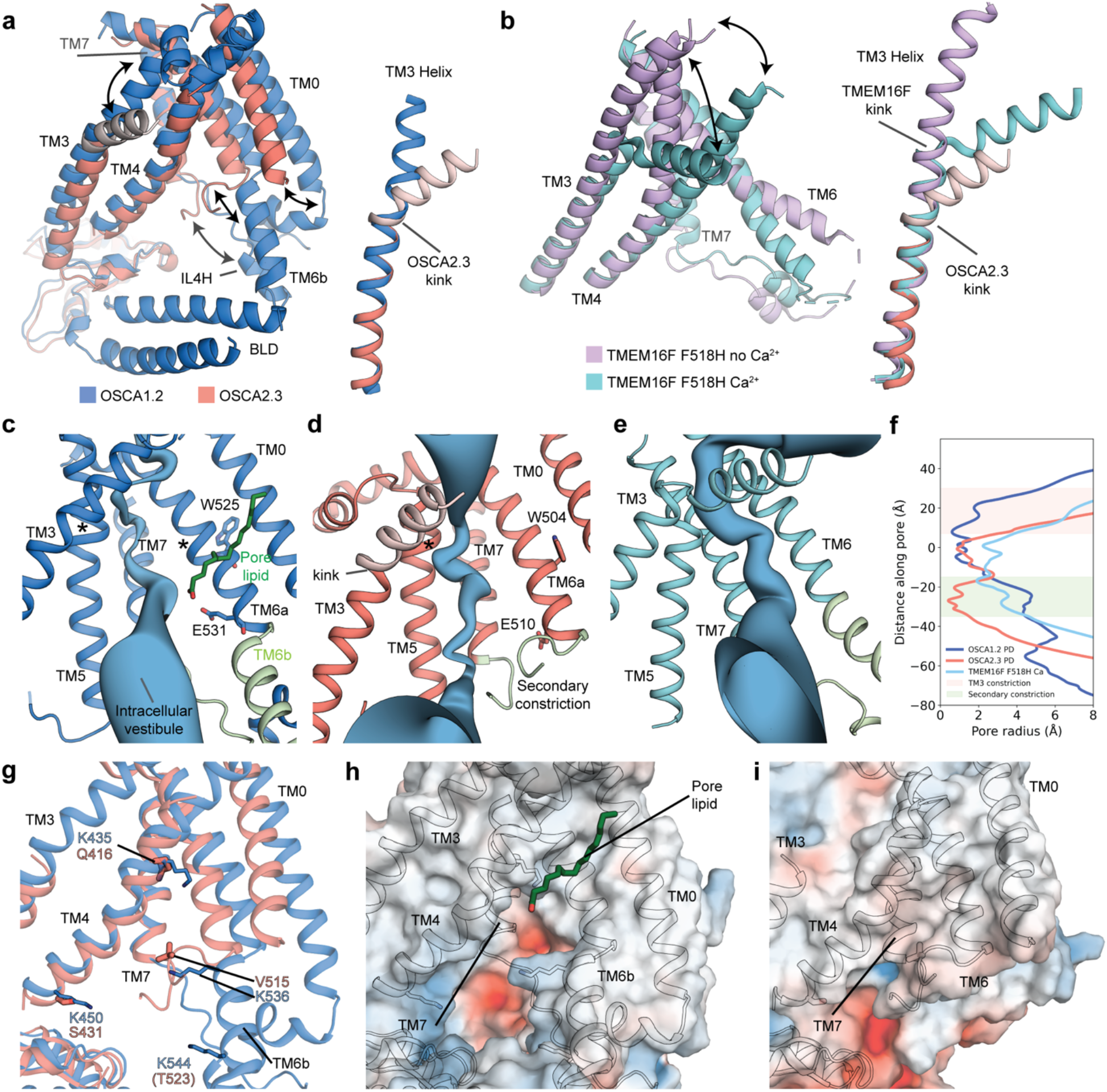
Pores and membrane fenestration. **a**, Superposition of pore lining TMs of OSCA1.2 and OSCA2.3 (left), with the movements of TM6a-TM7 cytoplasmic linker, TM0 and extracellular end of TM3 denoted by black arrows. Superposition of TM3 only (right), with kinked residues of OSCA2.3 colored light salmon. **b**, Superposition of pore lining TMs of TMEM16F F518H mutant structures with (PDB:8B8J) and without Ca^2+^ bound. (PDB:8B8J) (left), with the movements of the extracellular ends of TM3 and TM4 denoted by black arrows. Superposition of TM3 only of TMEM16F F518H mutants and OSCA2.3 (right). Depiction of pore pathway along the TMD of OSCA1.2 (**c**), OSCA2.3 (**d**) in peptidiscs and TMEM16F F518H Ca^2+^ bound (**e**). TM4 was omitted for clarity and the conserved glutamate residue of OSCAs and the π-helix tryptophan residue are shown as sticks and labeled. In green: pore lipid; light green: TM6a-TM7 linker and asterisks (**\***) denote π-helical turns. **f**, Pore radii profile of OSCA1.2 and OSCA2.3 in peptidiscs (PD), and TMEM16F F518H Ca^2+^ bound (PDB: 8B8J). Constriction caused by TM3 and secondary constriction is highlighted in light salmon and light green, respectively. **g**, Superposition of OSCA1.2 and OSCA2.3 showing predicted lipid-interacting lysine residues in OSCA1.2. These basic residues are not conserved in OSCA2.3. Electrostatic surface representation of OSCA1.2 (**h**) and OSCA2.3 (**i**). Extracellular end of TM3 of OSCA2.3 colored light salmon (**a, b, d**) is not included in (**i**) or the final model (see materials and methods).

Similar to TMEM16s^26-28^, the position of TM4 and TM6 of OSCA1.2 creates a fenestration in the TMD and make the pore accessible to the membrane lipids^20^ (Fig. 2g,h). Molecular dynamics (MD) simulations of OSCA1.2 (based on nanodisc structure) suggest lipids fill this opening and may block the pore pathway^20,21^. The presence of a lipid density in our OSCA1.2 structure in peptidisc supports the results of these simulations (Fig. 2h, Supplementary Fig. 2d). Furthermore, during the simulations, four basic residues located in TM4 and TM6b interacted tightly with the lipid phosphate group^20^. Interestingly, these four residues are not conserved in OSCA2.3, leading to a loss of positive charges in the area. TM0, TM6a, and TM6a-TM7 linker of OSCA2.3 are positioned closer to the intracellular end of the pore and seal the fenestration, preventing membrane lipids to access the pore (Fig. 2i).

### The BLD and IL4H are evolutionarily coupled

In OSCA1.2, the BLD and TM6b-IL4H are positioned adjacent to one other^20,22^ (Fig. 3a). Meanwhile in our OSCA2.3 map, density for the BLD was not observed and TM6b-IL4H has shifted away from the position of the BLD in OSCA1.2 (Fig. 1b, 2a). Thus, we asked whether these two structural changes were related. Coevolutionary sequence analysis performed on AtOSCA1.2 protein sequence revealed three pairs of coupled residues (probability > 0.86) between the BLD and IL4H (Fig. 3b,c). Despite these pairs not being in direct contact in the OSCA1.2 structure, closer inspection revealed that these sites favor residues capable of forming salt bridges: R/K257-E557, E260-R553, and R/K261-E/D554 (Fig. 3d). These results, along with the proximity of the residue pairs and the closeness of the BLD and IL4H, suggest that these two structural elements are evolutionary coupled. Interestingly, when the regions harboring these residue pairs are compared across the AtOSCAs, deletions of 2 of the pairs are present in clade 2 (Fig. 3e). Hence, we expect the interaction between the BLD and IL4H to be weakened in this clade, as seen in OSCA2.3 and OSCA2.2^21^.

**Figure 3.**
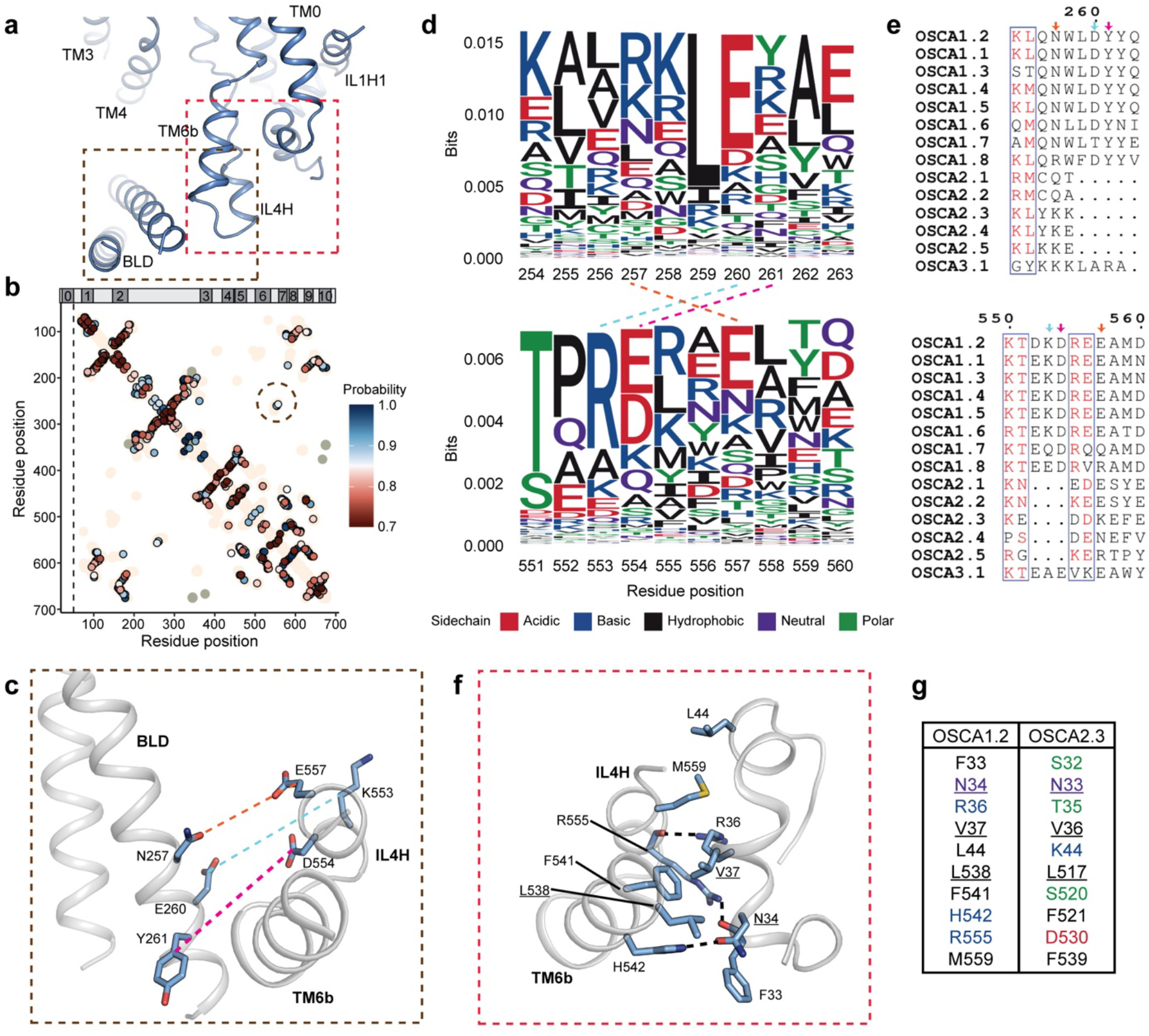
Interactions of the TM6a-TM7 linker with the BLD and TM0. **a**, Side view of OSCA1.2 centered on interactions between the BLD, TM6b/IL4H and cytoplasmic helix after TM0. **b**, Contact map from coevolutionary sequence analysis. Each points represents a pair of position predicted to be evolutionary coupled, colored based on probability of the coupling. Pairings closer than 6 positions in the sequence or have a probability of less than or equal to 0.7 are omitted. Tan and ash grey colored regions represent intra and inter subunit contacts, respectively, in experimental structures that are, on average, 5 Å or less apart. Circled are the coupled pairs between the BLD and IL4H. Bar on top represents the position of each TM in the sequence. **c**, Evolutionary coupled residues between BLD and IL4H depicted in OSCA1.2 structure. Sequence logo (**d**) and alignment (**e**) for the BLD (top) and IL4H (bottom) region that contains the coupled residues (circled in **b**). Dashed lines in **d** and arrows in **e** denote the coupled residues, color coded as in **c. f**, Interface between intracellular helix of TM0 and TM6b/ILH4. Dashed lines represent apparent hydrogen bonds. **g**, Conservation of residues in **f** between OSCA1.2 and OSCA2.3 based on sequence alignment. Residues are color coded as in **d**.

In OSCA1.2, TM6b-IL4H also contact the bottom helix of TM0 via hydrophobic interactions and hydrogen bonds (Fig. 3a,f). Yet, based on sequence alignment, only 3 out of 10 residues involved in this interface are conserved in OSCA2.3, and in most instances the properties of the side chains are altered (Fig. 3g). It is possible that disturbance of this interface further weakens the stability of TM6b-IL4H and thus they can shift to the position seen in OSCA2.3. Furthermore, this would free TM0 to move and explain its disposition away from the TMD core seen in our structure (Fig. 1b, 2d). Intriguingly, density corresponding to TM0 and TM6b-IL4H in OSCA2.2 structure in detergent^21^ resembles the disposition of the corresponding features of OSCA1.2 rather than OSCA2.3 (Supplementary Fig. 5a,b).

We used AlphaFold2^29^ (AF2) to predict the structures of monomers of OSCA2.2 and OSCA2.3 (OSCA2.2_pred_ and OSCA2.3_pred_, respectively) (Supplementary Fig. 5c,d). The disposition of the TMD of the AF2 predictions is closer to the peptidisc structure of OSCA1.2 than to the structure of OSCA2.3 and the TM6a-TM7 linker does not form the secondary constriction present on the intracellular side of the pore of the latter structure (Supplementary Fig. 5e). Furthermore, the predictions exhibit a low Predicted Alignment Error (PAE) for TM6b (residues 513-525) relative to two regions of interest. The first is the bottom of TM0 (residues 31-46), which TM6b contacts on the predicted structure. The second region corresponds to the intracellular ends of TM4 and TM5 (residues 424-445), which is proximal to TM6b in the OSCA2.3 structure and forms the secondary constriction (Supplementary Fig. 5f). The low PAE indicates a good prediction in the location of TM6b relative to these features. Additionally, OSCA2.2_pred_ provides a good fit in the OSCA2.2 experimental map solved in detergent^21^ (Supplementary Fig. 5c), suggesting the overall prediction for this protein is accurate. Likewise, if we assume a similar accuracy for OSCA2.3_pred_, these results suggest that OSCA2.3 may sample a conformation similar to the experimental structure of OSCA1.2 under the right set of conditions. We note that although AF2 does a remarkable job of structure prediction it does not capture the subtle conformational changes that are likely related to the function of OSCAs and as such there are many experimental structures yet to be solved to provide a molecular basis for mechanosensation in this family of proteins.

## Discussion

The coevolutionary sequence analysis performed in this study allows us to expand on previously proposed mechanisms of mechanosensation for OSCAs. It has been proposed that the membrane hook is capable of sensing membrane tension and displacing the BLD, which in turn can transmit the stimulus to the TMD to gate the channel^20-22^. Our analysis suggests that the BLD-IL4H interaction may play a role in linking the stimulus to the pore-lining helices TM6a and TM7. Whether the deletion of residues involved in these interactions affects mechanosensitive properties of clade 2 OSCAs, such as activation or inactivation kinetics, is yet to be determined as electrophysiological characterization of its members is lacking (with exception of AtOSCA2.3^7^). It is possible that charged residues around the deleted amino acids rescue these interactions to some extent. However, At clade 2 OSCAs (except for OSCA2.3) present other deletions near the membrane hook as well as the absence of the GXXGXXG motif that could debilitate the interaction between the BLD and the membrane and ultimately hinder mechanotransduction. Nonetheless, At clade 2 OSCAs present additional features, such as the amphipathic helix^20^, that could confer a mechanosensitive function to the channel.

Our cryo-EM structure of OSCA2.3 represents a previously unseen structural arrangement in the OSCA family. One possibility is that its structure represents a conformation specific to AtOSCA2.3. Notwithstanding, the similarity of the AF2 predictions and OSCA2.2^21^, which is closer in identity to OSCA2.3, to previously solved structures of clade 1^19-22^ and 3^19^ suggests that OSCA2.3 represents a new conformation accessible to other OSCAs. We hypothesize that the peptidisc environment used for structure determination could have been a crucial factor to obtain the conformation seen in OSCA2.3. While OSCA1.2 presents basic residues in the pore fenestration, these positive charges are not maintained in OSCA2.3, which likely decreases the strength of the interaction with lipids, resembling the effect of the R59L mutation in MscS^30,31^.

Tightly bound, annular lipids are expected to remain after detergent solubilization^32,33^, hence for OSCA2.3 weakly bound lipids would be replaced by detergent, resulting in a conformation similar to OSCA1.2 and OSCA2.2 structures in detergent^20,21^. However, detergent is removed at a later stage for peptidisc assembly^23^. The space left by the absence of lipids/detergent in OSCA2.3 is then filled by TM6a-TM7 linker to stabilize the channel, in a similar fashion to the rearrangement of the TMD of MscS after channel delipidation^31,34,35^. This hypothesis would not only effectively explain the different conformation seen between OSCA1.2 and OSCA2.3 in the same peptidisc environment, but it would also explain why the structure of OSCA2.2 in detergent does not present a similar conformation to OSCA2.3.

Previous MD simulations of OSCA1.2 suggested that lipid headgroups in the pore fenestration occlude the pore pathway on the intracellular side, thus delipidation may be necessary for the channel to become conductive^20^ (Fig. 4), similar to proposed mechanisms of TRAAK^36^ and MscS^31,34,35^. TM6 is a central component in the function of TMEM16 channels and scramblases^28,37,38^. Due the homology between TMEM16s and OSCAs, TM6 has been suggested to play a similarly important role in channel gating^19-22^.

**Figure 4.**
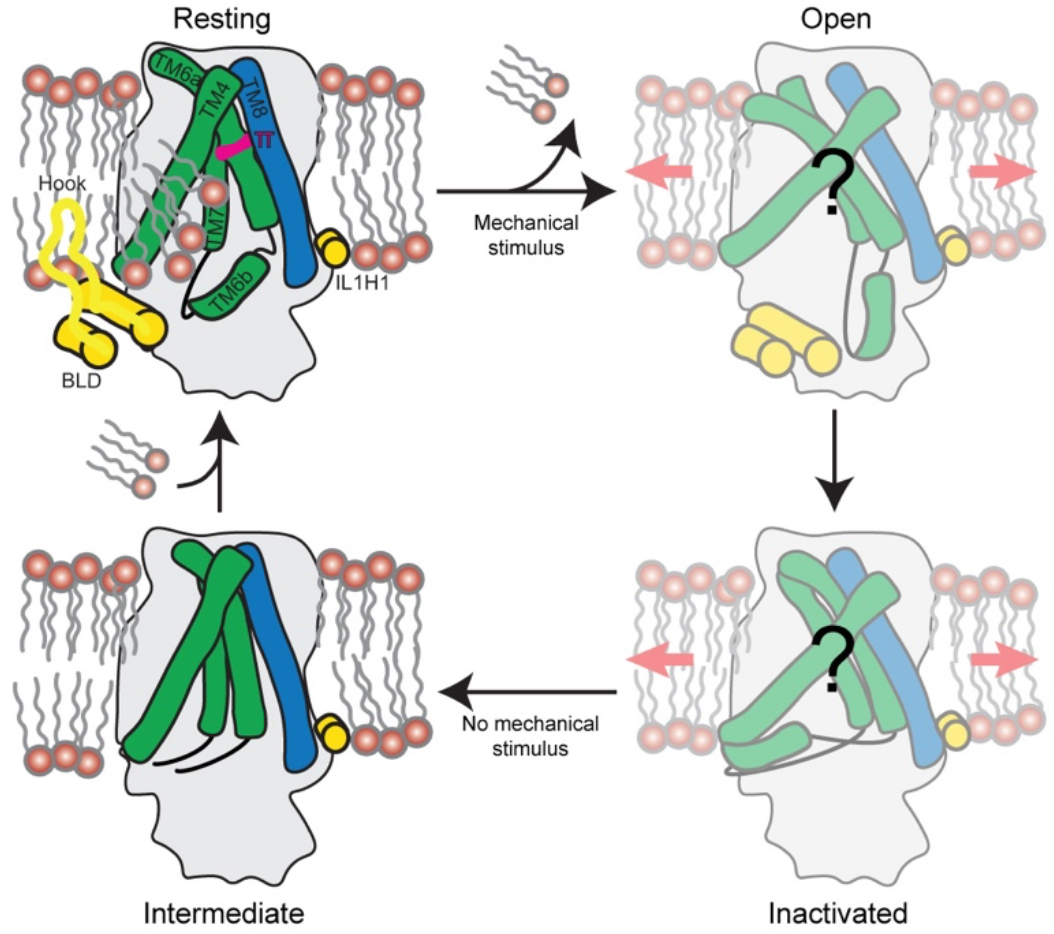
Proposed states of OSCA channels. In the absence of mechanical stimulus, the channel is in a resting state, with the hydrophobic gate closed and lipids populating the membrane fenestration. Upon membrane tension, the BLD or amphipathic helix (IL1H1) transmit the stimulus to the TMD, accompanied by delipidation of the pore pathway and the channel reorganizes into an unknown open state. Continuous stimulus causes the channel to inactivate, presumably by action of the TM6b-TM7 linker. Presumably, after the mechanical stimulus ceases, the hydrophobic gate closes, reaching the intermediate state represented by the OSCA2.3 structure in this study. Finally, the channel enters a resting state after lipids repopulate the pore fenestration.

Furthermore, MD simulations of OSCA1.1 under surface tension support the involvement of TM6 in gating, with mutagenesis and electrophysiology experiments advocating for a role of a conserved glycine as a potential hinge^19^. Therefore, a potential mechanism for gating of OSCA channels upon mechanical stimulus could involve exclusion of lipids from the pore pathway accompanied by shifting of TM6, possibly involving a π to α transition^20,22^ in a similar fashion to the α to π transition of the corresponding TM of mTMEM16A^28^, and resulting in an unknown conductive conformation. OSCAs exhibit inactivation when subjected to continuous stimulus^7^, but the mechanism through which it occurs is unknown. We speculate that, upon prolonged mechanical stimulus, TM6a-TM7 linker could move towards the pore fenestration and block the pore on the cytosolic side, like our structure of OSCA2.3, with the secondary constriction acting as an inactivation gate (Fig. 4). Given the hydrophobic gate is also closed in OSCA2.3, our structure could correspond to an intermediate state between the resting and inactivate conformations of the channel that we were able to capture due to lipid removal and absence of membrane tension. In this model, lipids would be required to repopulate the pore to return the channel to its resting state. Future work is needed to validate the existence of this proposed secondary gate in OSCAs and its role in inactivation.

Overall, our work reveals heterogeneity in the structural arrangement of the OSCA family, in particular the existence of EICs and motion of the TMD that altered the pore profile. Based on this novel conformation, we have updated our proposed model for mechanosensation in OSCA channels, which highlights the importance of lipids, as seen in other MA ion channels^31,34-36,39,40^. We also postulate the prospective existence of a secondary gate that limits ion flow upon persistence of stimulus, a feature proposed for other ion channels^41-47^, including MA Piezo channels^48-50^. We expect our work to serve as a foundation to further illuminate the molecular basis of mechanosensation and the structural diversity of the larger OSCA/TMEM63 family.

## Methods

### Expression constructs and viral production

Full-length human codon optimized OSCA2.3 (UniProt ID: Q8GUH7) was cloned into a pEG BacMam plasmid^51^ tagged at the C-terminus with a PreScission Protease cleavage site, followed by mGFP and Strep-tag II. A 4-5 residue linker was introduced between the gene and each one of the features mentioned. Bacmid DNA containing the construct of interest was transfected into 2mL of Sf9 cells at a density of 1×10^6^ cells/mL on a 6-well plate using ExpiFectamine Sf Transfection Reagent (Life Technologies) and following the manufacturer’s instructions. Cells were culture at 27ºC without shaking until the majority of cells exhibited strong GFP fluorescence (∼5-7 days). Supernatant was filtered and used to amplify virus for two subsequent generations. Baculovirus in the supernatant was pelleted and resuspended in FreeStyle 293 Expression Medium (Life Technologies) supplemented with 2% Fetal Bovine Serum (FBS). For OSCA1.2, a previously reported construct containing OSCA1.2-EGFP in pcDNA3.1 was used^20^.

### Protein expression and purification

For OSCA2.3, 3.6 L of Human Embryonic Kidney (HEK) 293F cells at 2.1×10^6^ cells/mL were transduced by addition of 10% v/v of baculovirus containing the construct of interest gene and incubated at 37ºC and 8% CO2 for ∼7 hours. 10mM sodium butyrate was added to the cells, followed by incubation at 32ºC for ∼45 h. Cells were pelleted by centrifugation, resuspended in ice-cold buffer A (TBS (25mM Tris pH = 8.0, 150mM NaCl), 2 μg/mL leupeptin, 2 μg/mL aprotinin, 1 mM phenylmethylsulfonyl fluoride (PMSF), 2 μM pepstatin A, and 2 mM DTT), and sonicated. Debris was pelleted by centrifugation and supernatant containing the cell membranes was pelleted for 45 min at 35000 rpm in a Type 45 Ti Fixed-Angle rotor. Membranes were stored at -80ºC for future use. Membranes were thawed on ice, then transferred and mechanically homogenized in a chilled Dounce homogenizer using buffer A. All subsequent steps were performed at 4ºC unless otherwise noted. Protein was then solubilized in buffer A supplemented with 1% n-dodecyl beta-D-maltopyranoside (DDM) and 0.1% cholesteryl hemisuccinate (CHS) for 1 hour with vigorous stirring. The insoluble fraction was pelleted using the Type 45 Ti Fixed-Angle rotor as outlined above. 2 mL of GFP nanobody^52^-coupled Sepharose resin made in-house was equilibrated with buffer W (TBS, 0.01% DDM, 0.001% CHS, 2 mM DTT). Batch binding of the supernatant to the resin was performed for 1 hour. Resin-bound protein was spun down at low speed and washed with buffer W twice and subsequently transferred into a gravity flow column, where it was washed again with approximately 10 CV of buffer W. Liquid was drained from the column, leaving just enough to cover the resin bed. 2 mg of NSPr peptide (Peptidisc Biotech) resuspended in ∼400 μL of ddH2O were added and the sample was transferred to a 14 mL conical using buffer P (TBS, 2 mM DTT), followed by a second addition of 2 mg of NSPr. We aimed for a molar ratio of 1:60 (protein:NSPr) or higher concentration of NSPr to provide an excess of peptide. The approximate OSCA2.3 yield was considered to be roughly the same as previous purifications where the protein was cleaved of the resin while still in detergent and absence of peptide. 125 mg of Bio-Beads SM-2 (Bio-Rad) (prewashed with methanol, twice with ddH2O and resuspended in TBS) were added and the mix was incubated for 1 h with rotation. 125 mg of Bio-Beads were added for a second time, along with 7 μL of PreScission Protease (Cytiva). Volume was adjusted to ∼10 mL and sample was incubated overnight under rotation.

The next day, the supernatant, resin, Bio-Beads mix was transferred into a gravity flow column and flow through containing the peptidisc-reconstituted sample was concentrated using a 100 kDA MWCO Amicon Ultra centrifugal filter (Millipore). Protein was then injected onto a Shimadzu HPLC for Size Exclusion Chromatography (SEC) using a Superose 6 increase (Cytiva) column. Fractions corresponding to OSCA2.3 reconstituted in peptidiscs were concentrated to ∼3.6 mg/mL (assuming 1 Absorbance Unit (AU) equal to 1 mg/mL) and used for preparation of cryo-EM grids on the same day.

For expression of OSCA1.2, 3.6 L of HEK293F cells were transfected and cultured as previously reported^20^. Membranes were obtained as outlined above and stored at -80ºC until needed. Purification and peptidisc reconstitution of OSCA1.2 was carried out similar to the OSCA2.3 sample, but with the changes described below. Buffer W was supplemented with 0.4 μM pepstatin A and 0.4 μg/mL aprotinin and it was used during the overnight incubation of the sample with Bio-Beads. The following day, the buffer W was replaced by buffer P using a gravity flow column, then returned to a conical tube and, after addition of ∼150 μg PreScission protease, the sample was incubated overnight. After SEC, the fractions corresponding to OSCA1.2 reconstituted in peptidiscs were concentrated to ∼8 mg/mL (assuming 1AU = 1 mg/mL).

### Cryo-EM sample preparation, data collection and processing

Cryo-EM grids were prepared using a Vitrobot Mark IV (ThermoFisher) operating at 4ºC and 100% humidity. 3.5 μL of sample (at ∼3.6 mg/mL for OSCA2.3 and ∼8 mg/mL for OSCA1.2) were applied to a plasma-cleaned UltrAuFoil 1.2/1.3 300 mesh grid. Grid was blotted once for 3.5-4 s after a wait time of 12 s and using with blot force 0, followed by plunge-freezing using nitrogen-cooled liquid ethane.

Data was collected on a Titan Krios (ThermoFisher) operating at 300 kV and using a K2 Summit direct electron detector (Gatan) with a pixel size and nominal magnification of 1.03 Å and 29,000x, respectively. Leginon^53^ was used for automated data collection, with 39 (OSCA2.3) and 36 (OSCA1.2) frames per movie for an accumulated dose of approximately 50 electrons per Å^2^. For OSCA2.3 sample, 5,323 micrographs across three grids were collected with a defocus range of -0.6 to -1.5 μm. For OSCA1.2, 1,805 micrographs with defocus range of -0.6 to -1.4 μm. The majority of movies were aligned and dose-weighted using MotionCor2^54^, while cryoSPARC^55^ full-frame motion correction was used for 88 micrographs of the OSCA2.3 due to availability of computational resources.

For OSCA2.3, dose-weighted micrographs were filtered using MicAssess^56^ with a threshold value of 0.05 and imported into cryoSPARCv2, followed by estimation of CTF values using Gctf^57^. Blob picker on the initial 608 micrographs returned 287,998 particles that were subjected to 2D classification followed by *ab initio* reconstruction.

The best out of two classes was further subjected to two subsequent rounds of homogeneous refinement (C1 and C2 symmetry imposed, respectively) and the final volume was used to generate templates. Templates were used to pick 2,683,410 particles from micrographs with CTF estimates better than 5 Å. 2D classification was used to sort out junk particles. *ab initio* reconstruction generated initial volumes that were subsequently used to further clean the particle stack into good and junk particles through heterogeneous refinement. 1,573,037 particles were subjected to homogeneous refinement without imposed symmetry and then the particle stack was imported into RELION^58^. The particle stack underwent, in order, 3D refinement, 3D classification without alignment (T = 4), 3D refinement, introduction of optics groups based on which grid each micrograph came from, CTF refinement and 3D refinement. These 3D refinements did not imposed symmetry and only allowed for local angular searches. At this point, we decided to return to the raw movies in order to perform Bayesian polishing^59^. Movies were aligned and dose weighted using the RELION implementation of MotionCor2^54,59^, followed by CTF estimation with CTFFIND4^60^. Due to frames not being rotated equally between the two motion correction stages (or flipped equally for the case of the movies aligned in cryoSPARC), the coordinates for each particle were converted to match the new rotation outside of RELION with an in-house script, followed by a 3D refinement imposing C2 symmetry and with global searches to realign the particles. After this point, unless otherwise specified, all 3D refinements impose C2 symmetry, only allow local searches, and use SIDESPLITTER^61^ for the reconstruction step. 3D classification (T = 8) produced one class (322,432 particles) that resembled an OSCA dimer, which was then subjected to 3D refinement (C1), and one round of CTF refinement and polishing with 3D refinements after each. A final 3D classification without alignment (T = 20) returned a final stack of 180,640 particles, that were CTF refined and polished (with a 3D refinement after each step) for 2 more rounds.

For OSCA1.2, dose-weighted micrographs were imported directly into cryoSPARC followed by Gctf. A subset of 250 micrographs was used to pick 82,586 blobs, which were subjected to two rounds of 2D classification and *ab initio* reconstruction. Non-uniform (NU) refinement, global and local CTF refinement, and two rounds of NU refinement were done on the best class. Only the last NU refinement imposed C2 symmetry. The resulting volume was used to generate templates, which were later used to pick 1,046,049 particles on the full dataset, followed by 2D classification and *ab initio* reconstruction. The 2 best classes were subjected to NU refinement, separately, and then combined for a heterogeneous refinement. The particle stack of the best class, consisting of 315,944 particles, was used for NU refinement before being imported into RELION. An initial 3D refinement (global angular searches) was followed by 3D classification without alignment (T = 20). 3D refinements in RELION only allowed for local angular searches and did not imposed symmetry unless specified otherwise.

Particles belonging to the best class underwent the following steps, in order: 3D refinement, and 3 rounds of CTF refinement with 3D refinements in between. To estimate the correct symmetry axis, one 3D refinement with C2 symmetry and global angular searches was performed, and after this point the remaining refinements imposed this symmetry. The particle stack was subjected to a 3D refinement, a fourth CTF refinement and 3D classification without alignment (T = 20). A final 3D refinement was performed on the best class (57,250 particles).

Each final map was postprocessed with LocalDeblur^62^ using Scipion^63-65^ and, separately, DeepEMhancer^66^ using the ‘highRes’ model. The FSCs and local resolutions of the maps were calculated with RELION.

### Model building and refinement

A homology model of OSCA2.3 was obtained using SWISS-MODEL^67^ using the structure of OSCA1.2 in nanodiscs^20^ (PDB ID: 6MGV) as a template. This same template was used as the starting model for OSCA1.2 in peptidisc. Models were fit into the corresponding density using UCSF Chimera^68^, followed by visual inspection and building in Coot^69,70^. ELBOW^71^ was used to obtain the restrains from the different ligands. The final models were obtained through iterative rounds of building and real space refinement in Coot, Phenix^72,73^ and Rosetta^74^. ISOLDE^75^ was used to model TM3 of OSCA2.3 due to low quality of the map in this region. Molprobity^76^, EMringer^77^ and Phenix mtriage^78^ were used to validate the models against the appropriate LocalDeblur map. For OSCA2.3, the final model includes residues 3-26, 65-115, 129-225, 302-368, 410-517 and 536-691. For OSCA1.2, the model includes residues 2-50, 71-121, 157-266, 290-487, 505-648, 654-718. The pore profiles were obtained using the program CHAP^79^. Due to low resolution of TM3 of OSCA2.3, the extracellular portion of this TM could not be built confidently, which artificially deviated the direction of the pore pathway. To ameliorate this effect, the structure of OSCA2.3 was predicted with AF2^29^ and prediction and experimental model were aligned on TM3. Due to the good overlap between the two TMs, the extracellular end of AF2 prediction was grafted onto the experimental model, which included residues 368-381. Subsequently, Coot was used to fit this stretch of residues into the corresponding density of an unsharpened OSCA2.3 map gaussian-pass filtered to 2 σ. This model was used for pore profile calculation with CHAP and for figures (TM3 graft is colored light salmon), but residues 369-381 are absent from the deposited model. To calculate the cross-sectional area, distances between a residue and its counterpart on the complementary protomer were measured in Chimera. These distances were considered diagonals of a rhombus and the area was calculated accordingly. The residues used for OSCA2.3 (and complementary residues for OSCA1.2) were: W455 (V474) and P635 (A660) for the top measurement, and W353 (T372) and I601 (D626) for the bottom.

Structure figures were made using PyMOL^80^, UCSF Chimera and UCSF ChimeraX^81^. Clustal omega^82^ was used to reproduce the amino acid sequence alignment from a previous publication^20^. ESPript3^83^ was used to displayed only the sequences from *Arabidopsis thaliana* that have an available model (Supplementary Fig. 3) or the sequences from clades 1, 2 and 3 (Fig. 2g).

### Coevolutionary sequence analysis

Coevolutionary sequence analysis was performed using the EVcouplings software^84^ (https://evcouplings2.hms.harvard.edu/) with default inputs. Residues 1-700 of *A. thaliana* OSCA1.2 sequence (UniProt ID: Q5XEZ5) were used as input for the analysis. A total of 5439 sequence were used for the analysis with a bitscore threshold of 0.1 and 5 search iterations against the UniRef90 database. The resulting residue frequency per site was used to plot the sequence logos using the ggseqlogo^85^ R package.

## Data availability

Atomic coordinates for models of OSCA2.3 and OSCA1.2 in peptidiscs have been deposited to the Protein Data Bank (PDB) under IDs xxxx and xxxx, respectively. Corresponding cryo-EM maps have been deposited Electron Microscopy Data Bank (EMDB) under accession numbers yyyyy (OSCA2.3) and yyyyy (OSCA1.2), with the LocalDeblur map as the primary map.

## Author contributions

S.J.C. and W.H.L. cloned expression constructs. S.J.C., B.B. and W.H.L. prepared protein samples. S.J.C. and B.B. acquired cryo-EM data, processed data, and built and refined the atomic structures. A.B.W. supervised the project. S.J.C. and B.B. drafted the manuscript, which was edited and finalized with contributions from all authors.

## Acknowledgements

We thank W. Anderson for managing the electron microscopy facility at Scripps Research, J. Torres for helping with data collection, and C. Bowman, L. Dong and J.C. Ducom for assistance with computation. We acknowledge members of the Ward lab and M. Mravic for helpful advice. We thank S. Murthy for valuable discussion and feedback on the manuscript. This work was supported by NIH grant R01 HL143297 and a Ray Thomas Edwards Foundation grant to A.B.W.

Molecular graphics and analyses performed with UCSF Chimera and UCSF ChimeraX, developed by the Resource for Biocomputing, Visualization, and Informatics at the University of California, San Francisco, with support from National Institutes of Health R01-GM129325 and P41-GM103311, and the Office of Cyber Infrastructure and Computational Biology, National Institute of Allergy and Infectious Diseases.

