## Supplemental Figures for "Structure of mechanically activated ion channel OSCA2.3 reveals mobile elements in the transmembrane domain"

### Supplementary Figures:

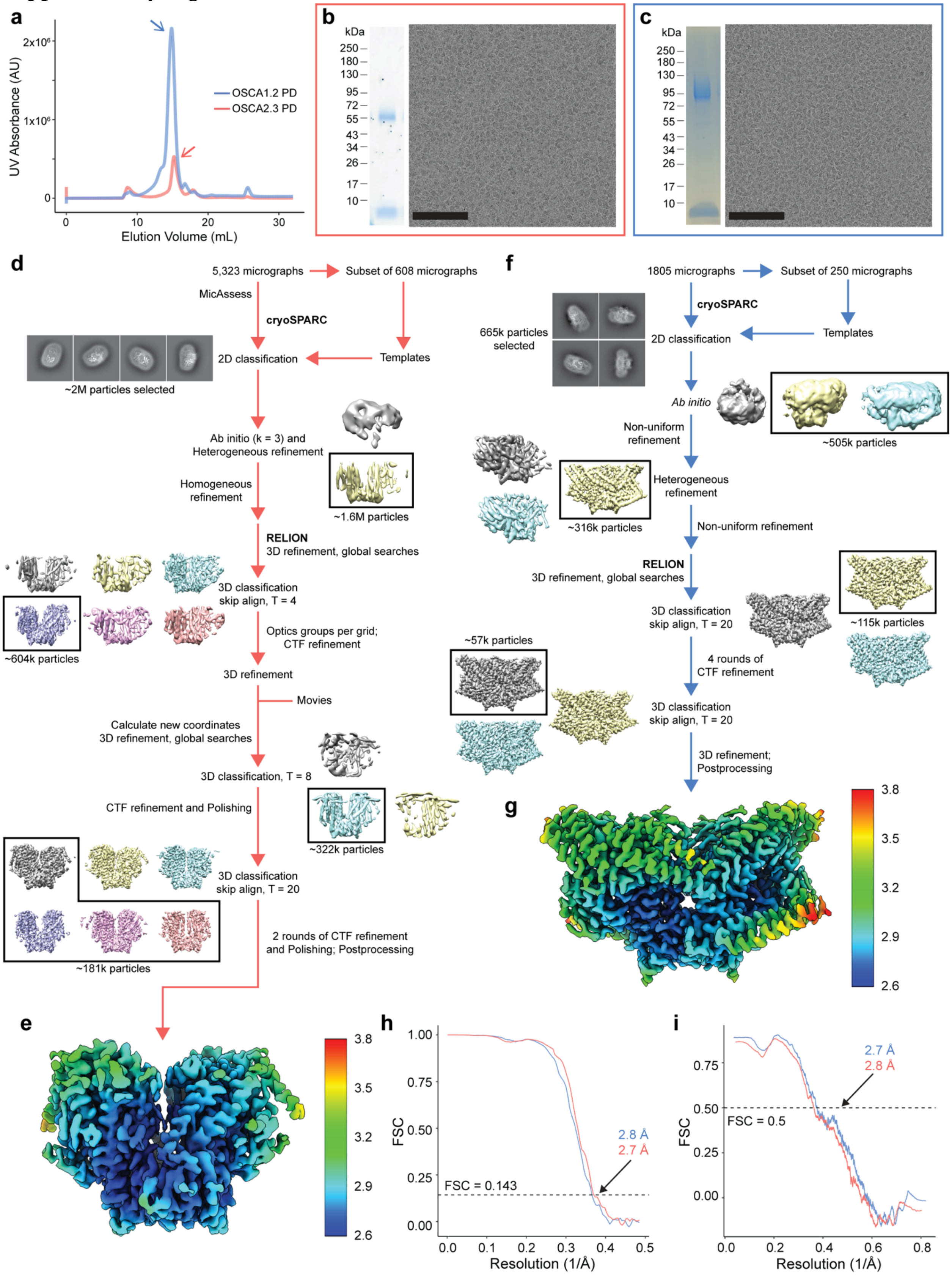

**Supplementary Figure 1. Purification and cryo-EM data processing of OSCA2.3 and OSCA1.2 in peptidiscs.**

**a,** SEC trace from OSCA2.3 and OSCA1.2 reconstitution in peptidiscs. Arrow points to peak corresponding to fractions pooled and used for cryo-EM. **b,** Left: SDS-PAGE of pooled fractions containing OSCA2.3 reconstituted in peptidiscs. Molecular weight of OSCA2.3 and peptidisc peptide is approximately 80 and 4.5 kDa, respectively. Right: Representative cryo-EM micrograph. Black bar is 100nm. **c,** SDS-PAGE (left) and representative micrograph (right) for OSCA1.2 in peptidiscs sample. Molecular weight of OSCA1.2 is approximately 88 kDa. **d,** Cryo-EM processing workflow for OSCA2.3 dataset. **e,** OSCA2.3 LocalDeblur map colored by local resolution calculated in RELION. **f,** Cryo-EM processing workflow for OSCA1.2 dataset. **g,** OSCA1.2 LocalDeblur map colored by local resolution calculated in RELION. **h,** FSC plot calculated in RELION. **i,** LocalDeblur map to model FSC plot.

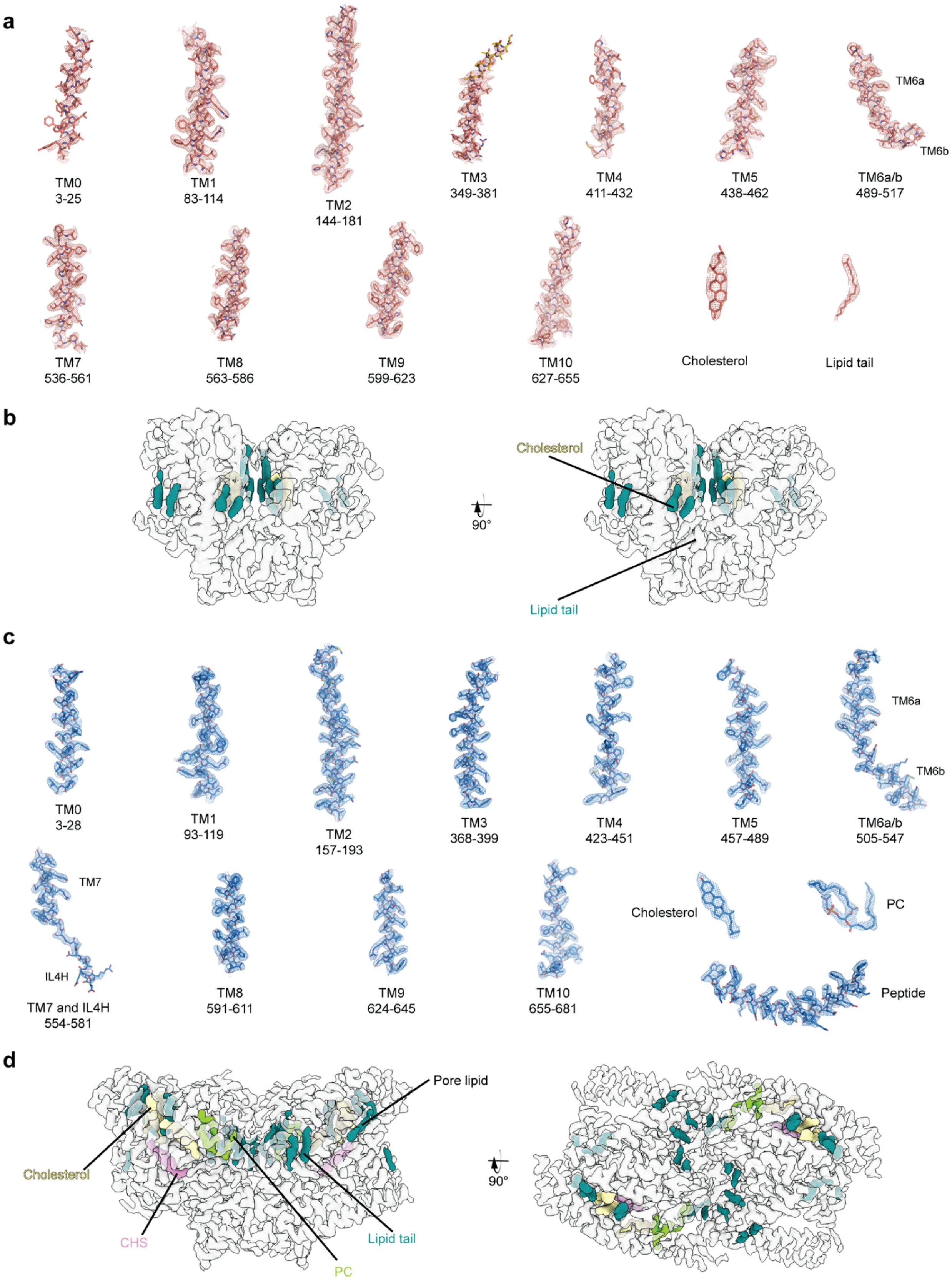
a
TM0
3-25
TM1
83-114
TM2
144-181
TM3
349-381
TM4
411-432
TM5
438-462
TM6a
TM6b
TM6a/b
489-517
TM7
536-561
TM8
563-586
TM9
599-623
TM10
627-655
Cholesterol
Lipid tail
b
Cholesterol
90°
Lipid tail
c
TM0
3-28
TM1
93-119
TM2
157-193
TM3
368-399
TM4
423-451
TM5
457-489
TM6a
TM6b
TM6a/b
505-547
TM7
IL4H
TM7 and IL4H
554-581
TM8
591-611
TM9
624-645
TM10
655-681
Cholesterol
PC
Peptide
d
Pore lipid
Cholesterol
CHS
PC
Lipid tail
90°

**Supplementary Figure 2. Fit of OSCA2.3 and OSCA1.2 models to respective LocalDeblur map.**

Fit of selected regions of the OSCA2.3 (**a**) and OSCA1.2 (**c**) models to the respective density in the LocalDeblur map. Maps were contoured at a threshold of 5 σ. For OSCA2.3 TM3, residues grafted from the AlphaFold2 (AF2) model of OSCA2.3 are colored in yellow and are modeled as poly-alanine. Highlight of densities in the OSCA2.3 (**b**) and OSCA1.2 (**d**) maps that are modeled as lipids. CHS: cholesteryl hemisuccinate; PC: phosphatidylcholine. Densities labeled as lipid tails were modeled as palmitic acid.

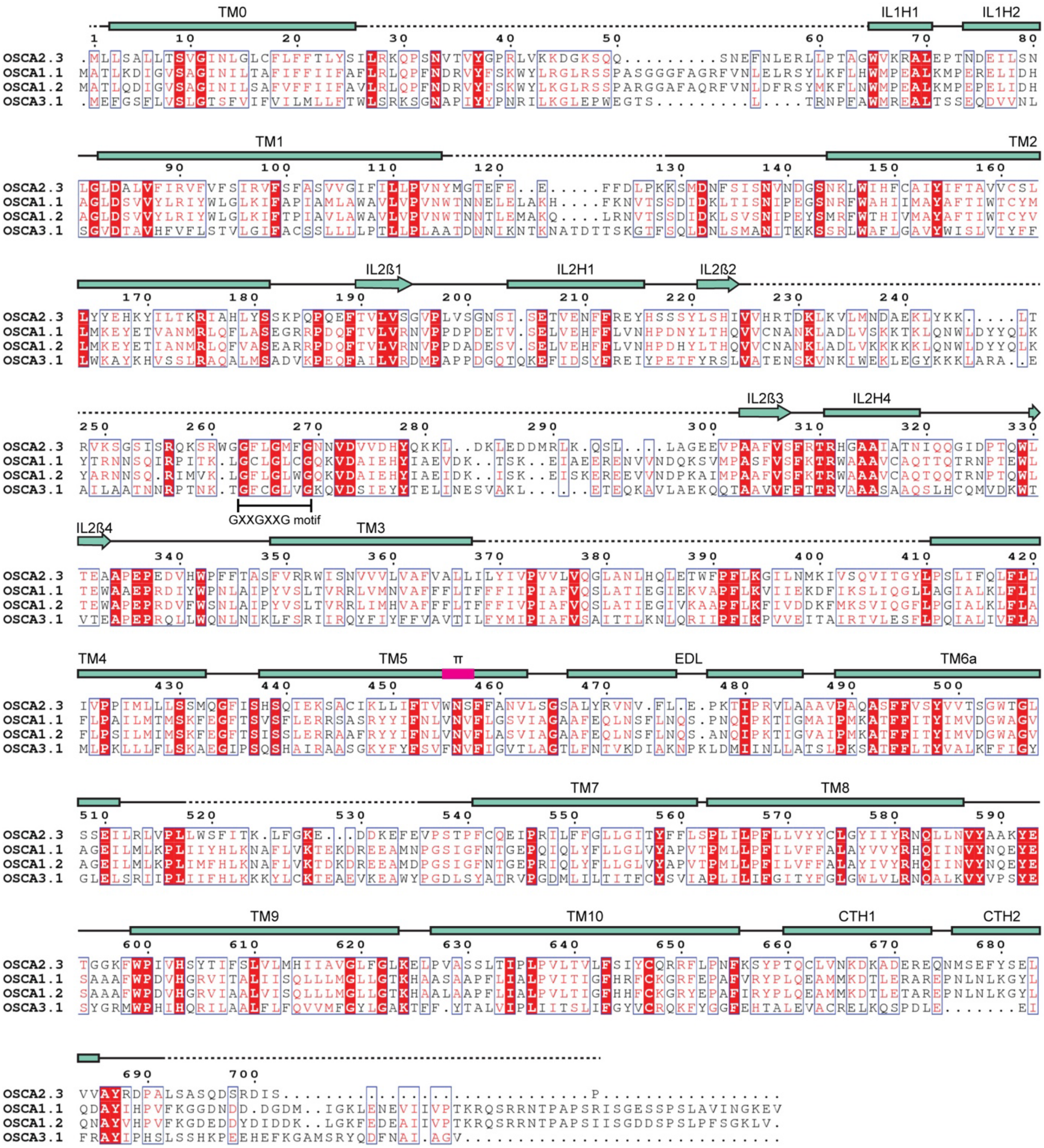


**Supplementary Figure 3. Amino acid sequence alignment of OSCA2.3.**

Amino acid sequence alignment of *Arabidopsis thaliana* OSCA2.3, OSCA1.1, OSCA1.2 and OSCA3.1. Secondary structure of OSCA2.3 represented on top, with helices as rectangles and ß-strands as arrows. Dashed lines correspond to regions not modelled. π helical turn labeled in magenta. TM: Transmembrane; IL: Intracellular Loop; CT: C-terminal; H: Helix; ß: ß-sheet; EDL: Extracellular Dimerization Loop. The BLD, formed by IL2H2 and IL2H3, is absent as these helices were not observed in the OSCA2.3 map.

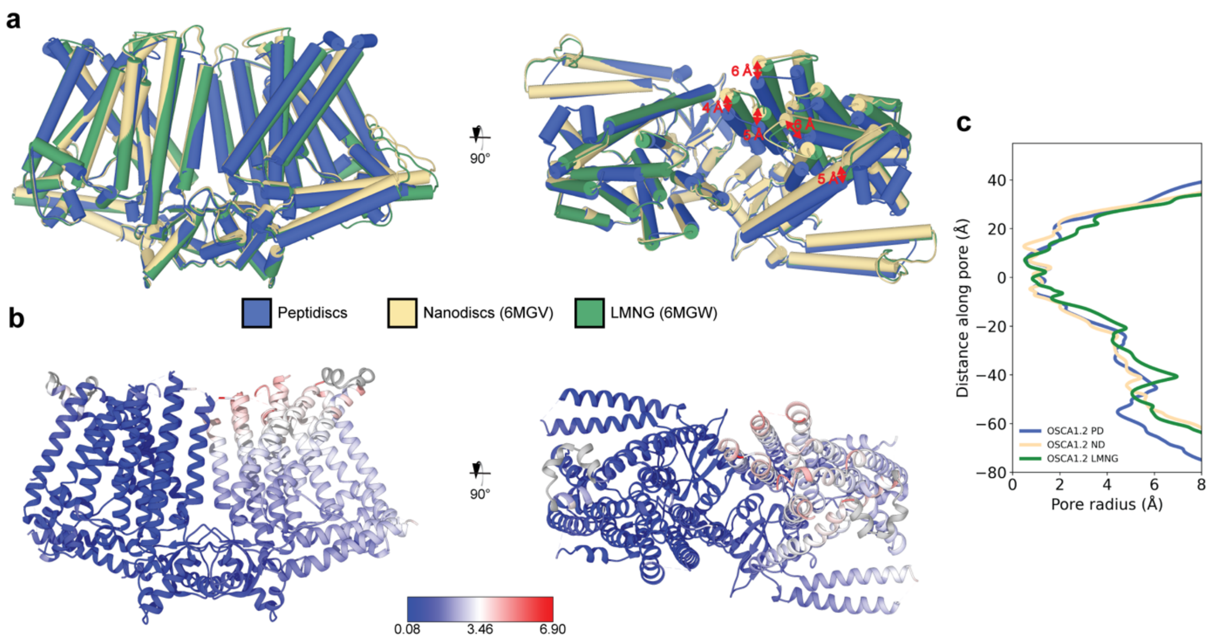


**Supplementary Figure 4. Comparison of OSCA1.2 structures solved in different environments.**
**a,** Superposition of OSCA1.2 structures in peptidiscs, nanodiscs and detergent (LMNG) aligned to left protomer. Movement between helices of right protomer is measured between carbon alpha of residues near end of helix between peptidisc and nanodisc structures. **b,** OSCA1.2 in peptidisc colored by RMSD (in Å) relative to LMNG and nanodisc structures. **c,** Pore radii profiles of OSCA1.2 in peptidisc (PD), nanodisc (ND) and LMNG.

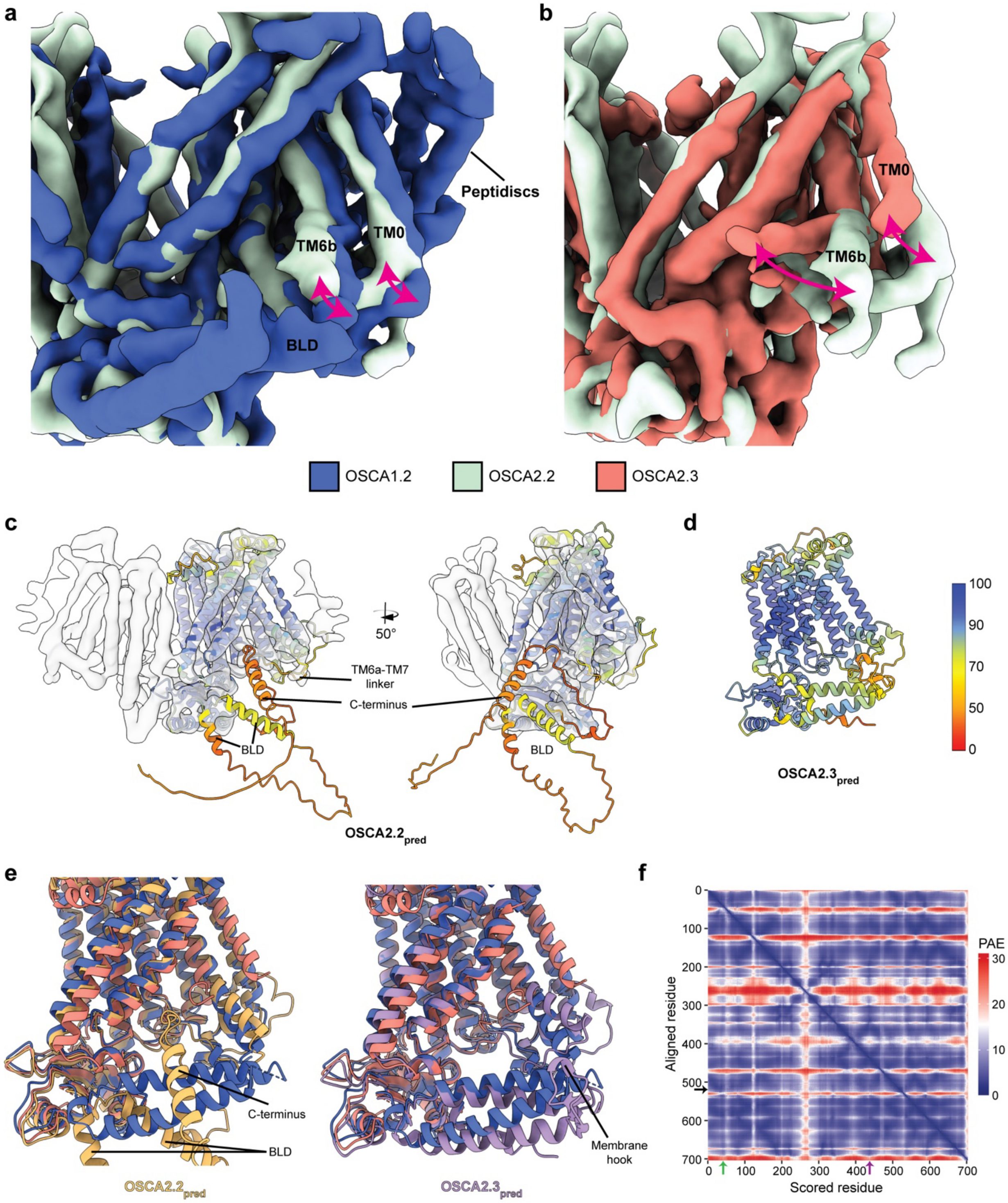


**Supplementary Figure 5. Comparison of OSCA2.2 map (EMDB: 9677) relative to OSCA1.2 and OSCA2.3.**

**a,** Superposition of OSCA1.2 and OSCA2.2 maps showing little displacement of TM0 and TM6b but absence of BLD in OSCA2.2. **b,** Superposition of OSCA2.2 and OSCA2.3 maps showing displacement of TM0 and TM6b and absence of BLD in both maps. Alignment was done on the protomers shown. Maps of OSCA1.2 and OSCA2.3 correspond to the unsharpened maps low pass filtered to 5.4 Å. **c,** Fit of AF2 prediction for OSCA2.2 to cryo-EM map of the same protein in detergent (EMDB: 9677). **d,** AF2 prediction for OSCA2.3. AF2 predictions in **c** and **d** are colored by predicted local-distance difference test (pLDDT) confidence measure, with higher values representing regions of higher local model quality. **e,** Superposition of structures of OSCA1.2 and OSCA2.3 to AF2 predictions of OSCA2.2 (left) and OSCA2.3 (right). **f,** PAE matrix for OSCA2.$3_{pred}$. Arrows point to the general area of TM6b (black, residues 513-525), bottom of TM0 (green, residues 31-46), and intracellular ends of TM4 and TM5 (purple, residues 424-445).
